# DeepTAP: an RNN-based method of TAP-binding peptide prediction in the selection of tumor neoantigens

**DOI:** 10.1101/2023.02.13.528393

**Authors:** Xue Zhang, Jingcheng Wu, Joseph Baeza, Katie Gu, Zhan Zhou

**Affiliations:** Innovation Institute for Artificial Intelligence in Medicine, College of Pharmaceutical Sciences, Zhejiang University, Hangzhou 310058, China; Biology Program, Iowa State University, Ames, IA 50011, USA

**Author notes:** Volunteer student from Ames High School, Ames, Iowa, USA. E-mail address (Katie Gu). To whom correspondence should be addressed. E-mail addresses (Z. Zhou), (J. Baeza).

## Abstract

The transport of antigenic peptides from cytoplasm to the endoplasmic reticulum (ER) via transporter associated with antigen processing (TAP) is a critical step during the presentation of tumor neoantigens. The application of computational approaches significantly speed up the analysis of this biological process. Here, we present a tool named DeepTAP for TAP-binding peptide prediction, which employs a sequence-based multilayered recurrent neural network (RNN). Compared with traditional machine learning and other available prediction tools, DeepTAP achieves state-of-the-art performance on the benchmark datasets. The source code and dataset of DeepTAP are available freely via GitHub at https://github.com/zjupgx/DeepTAP.

## Introduction

Cytotoxic (CD8+) T cells can recognize the peptides presented by major histocompatibility (MHC) class I molecules on the cell surface. These peptides are usually generated from the degradation of endogenous proteins, especially pathogenic or tumor-related proteins (Rossjohn *et al.*, 2015). The presentation of tumor neoantigen is produced by a series of complex intracellular processes. For example, proteins in proteasome are degraded into peptides, further cleaved by cytoplasmic peptidase, transported by the transporter associated with antigen processing (TAP) to ER, modified by ER-resident aminopeptidase (ERAP), and stably linked to one of several MHC class I molecules, finally presented on the cell surface (Kotsias *et al.*, 2019).

Bioinformatics and computational methods have been widely used to predict such epitopes processing and presentation. Although peptides binding to MHC class I is the most selective step, the translocation of peptides into the ER is a prerequisite, which limits or regulates the supply and binding of peptides for MHC class I molecules and, ultimately, immunogenicity (Sette *et al.*, 1994; Zhong *et al.*, 2003). Several tools have been developed to predict TAP-binding peptides, based on support vector machine (SVM) such as TAPPred (Bhasin and Raghava, 2004), TAPREG (Diez-Rivero *et al.*, 2010), or scoring matrix methods such as SMM (Peters *et al.*, 2003), KSMM (Hao *et al.*, 2020), or artificial neural network such as PREDTAP (Zhang *et al.*, 2006), as well as gaussian process regression (Ren *et al.*, 2011). However, the accuracy of these tools still needs to be improved, and the emergence and rapid development of new deep learning methods provide an opportunity for accurate prediction of TAP binding. In this study, we expand the TAP-binding peptide dataset and propose an RNN-based tool DeepTAP, which has been shown to predict TAP binding peptides accurately and outperformed other acknowledged tools on the benchmark dataset.

## Implementation

Currently, the public available TAP-binding dataset was compiled by Diez-Rivero, which contains 613 pieces of data (DS613), the label is log (relative IC_50_) referenced to RRYNASTEL (Diez-Rivero *et al.*, 2010). Here, we used this dataset to train regressor, and also collected several new binding data from the MHCBN database and related literature, collating a larger dataset (DS1114) consisting of 1114 items for classification training. Gated recurrent units (GRU) and long short-term memory (LSTM), further combined with bidirectional as well attention mechanisms, are considered as the main framework of the model, employing five-fold cross-validation for model training on classification and regression datasets respectively. Results show that the bidirectional gated recurrent unit (BGRU) achieves the best prediction performance (Table S1), which is used as the architecture of our model. DeepTAP is implemented based on Python3.9.7 and Pytorch1.9.1. The compiled dataset and code are open source and can be implemented, also with the usage documents are available freely on GitHub.

## Results

We first organized all collected TAP binding peptides, finding that they have a similar sequence length of 8-15mer, and the vast majority of 9mer peptides account for more than 80%. In more detail, TAP has a transport preference for the aromatic hydrophobic residues at the C-terminal, the charged hydrophobic residues in p2, and the hydrophobic residues in p3. Then, the architecture of DeepTAP is as shown in Fig. 1 a, briefly, the classifier and regressor were trained using DS1114 and DS613 respectively, taking as input a variable-length 8-15mer peptide, followed by embedding and three RNN hidden layers. Classifier outputs a prediction score ranging from 0 to 1, above 0.5 considered to be a positive one, with regards to regressor is the log value of relative IC_50_.

**Fig. 1.**
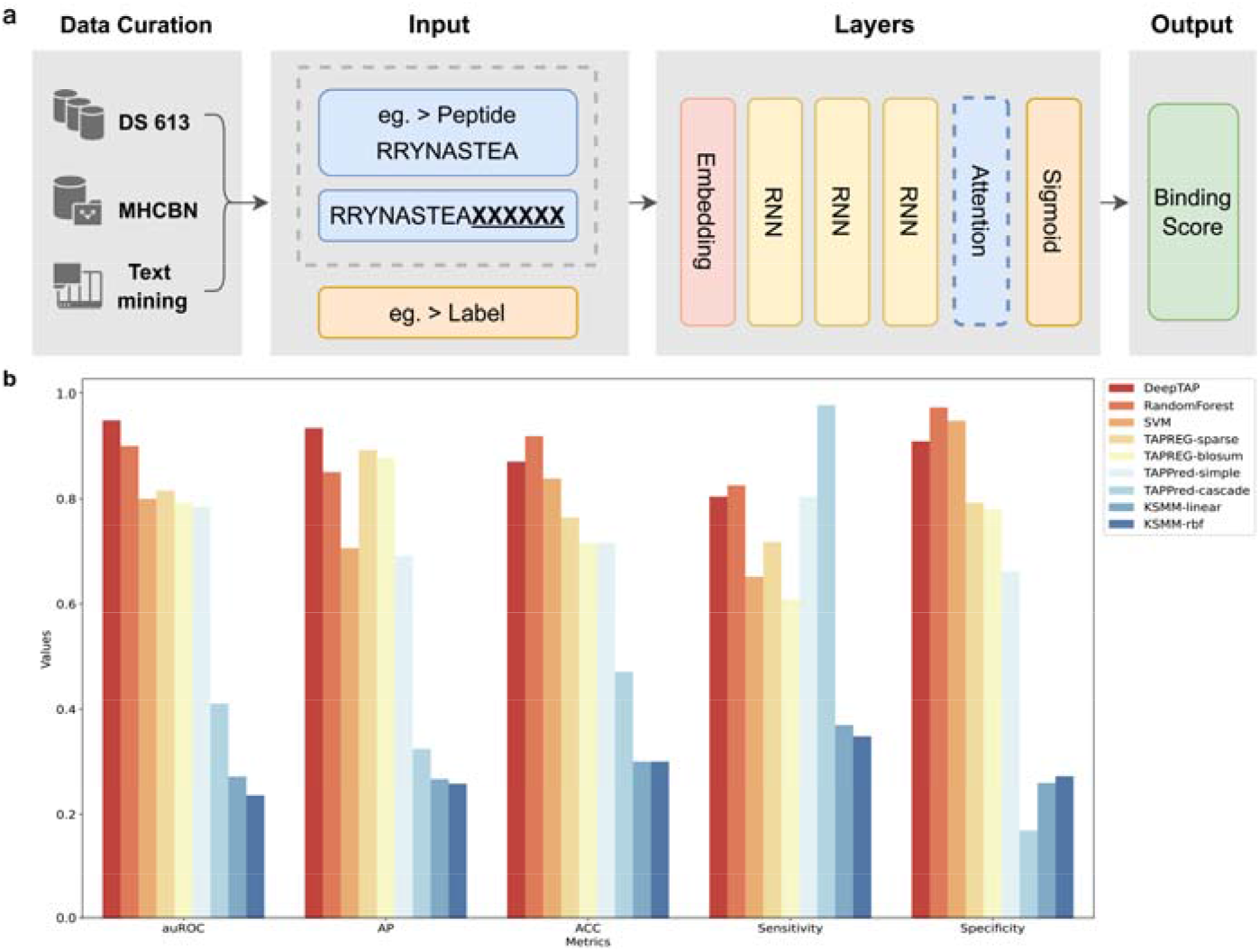
The schematization and performance of DeepTAP. The architecture of DeepTAP with input data, embedding, hidden layers and output are shown in (a) and classification model performances compared with other tools are displayed in (b).

To evaluate the prediction performance, we complied a dataset containing 120 pieces of TAP-binding peptides, not used in the training process and previous tools, used to benchmark our classification model and other baselines, including traditional machine learning methods, support vector machine (SVM) and random forest (RF), as well as available prediction tools, TAPPRED, TAPREG as well KSMM. For the regression tasks, it cannot be directly compared with the existing tools, the evaluation results of each tool are directly quoted from the paper. Among them, TAPPRED consists of two patterns, a simple SVM and a two-layer cascade SVM, TAPREG is based on traditional SVM and KSMM is a matrix algorithm considering the pairwise interaction between amino acids.

DeepTAP achieved the best prediction result on the classification benchmark dataset, with AUC reaching 0.95, followed by RF with an AUC of 0.90. Meanwhile, the AUC values of several tools based on SVM are relatively consistent, around 0.8. However, cascade SVM and KSMM are evaluated lower than 0.5, which is worse than the random prediction (Fig. 1b, Table S2, Fig. S1, Fig. S2). Parallelly, our model also achieves state-of-art performance in regression tasks with the Spearman correlation coefficient reaching 0.9 (Table S3). Overall, the results strongly indicated that compared with other prediction tools DeepTAP achieved higher accuracy and confidence prediction, and showed significant advantages in the TAP-binding peptide prediction task.

## Conclusions

In this study, we compiled a TAP-binding peptide dataset and report DeepTAP, an RNN-based tool for TAP-binding peptide prediction, which has been evaluated and compared with traditional machine learning SVM, RF and other available tools. The results show that DeepTAP is superior to the existing prediction tools in terms of prediction accuracy and error control. As far as we know, DeepTAP is the first application of the deep learning method on TAP-peptide binding prediction. To ensure easy installation and usage, as well as high portability, reproducibility and scalability, the DeepTAP tool is available on GitHub (https://github.com/ziupgx/DeepTAP).

## Acknowledgements

This work was supported by the National Natural Science Foundation of China [Grant No. 31971371]. We thank the Information Technology Center and State Key Lab of CAD&CG Zhejiang University, and Alibaba Cloud for the support of computing resources.

