## Supplemental information for "DeepTAP: an RNN-based method of TAP-binding peptide prediction in the selection of tumor neoantigens"

**To whom correspondence should be addressed.*

### Supplementary Methods

#### Datasets

The data of TAP-binding peptides used in the study were collected from the MHCBN database and also other relevant literature. The sources of data are as follows: (1)Up to March 2022, there are 1053 TAP binding sequences in the MHCBN database(Bhasin *et al.*, 2003). Among them, 643 peptides were bound to human TAP, and 410 binding data were obtained by deleting redundant and incorrect sequences. (2) Dize-Rivero et al. compiled a dataset containing 613 peptides binding to human TAP (DS613), incorporating 435 peptides provided by Daniel et al. and 178 peptides from the AntiJen database(Daniel *et al.*, 1998; Diez-Rivero *et al.*, 2010; Toseland *et al.*, 2005). (3) In addition, text mining was employed to search the related literature, and 163 TAP binding peptides were fortunately obtained from nine articles (Androlewicz and Cresswell, 1994; Armandola *et al.*, 1996; van Endert *et al.*, 1994; Gorbulev *et al.*, 2001; Koopmann *et al.*, 1996; Momburg *et al.*, 1994; Neefjes *et al.*, 1995; Obst *et al.*, 1995; Shepherd *et al.*, 1993). The data from the above three ways were merged and sorted out, deleting duplicated sequences and a few sequences exceeding 15mer (less than 7). Moreover, 120 pieces of data were compiled as an independent benchmark dataset. Ultimately a dataset containing 1114 TAP-binding peptides (DS1114) and a benchmark dataset (DS120) were created for the training and evaluation of the classification model. The dataset DS613 was used for regressor training, which was labelled as the log value of relative IC50 referenced to RRYNASTEL concerning each peptide.

#### Encoding method

The machine learning algorithm cannot analyze the amino acid sequence directly, so it is necessary to convert amino acids into numbers that can be recognized before they can be used as input. Sparse coding is used in this study, also known as one-hot coding, mainly uses N-bit state registers to encode N states, each state has an independent register bit, and only one bit is valid at any time. Owing to its simplicity and effectiveness, this method is widely used in available prediction tools. In this study, considering padding sequence to 15mer with pseudo-amino acid "X", each amino acid is converted into a unique vector with twenty zeros and a single one, and finally, each peptide sequence is represented as a matrix of $n \times 21$.

**Model selection**

A recurrent neural network (RNN) is acknowledged as a popular theoretical framework for supervised generalization, which has evident advantages over other networks in extracting temporal and semantic information embedded in data. The early RNN is relatively simple in that the input received by the neuron node at a certain time includes the input of the current time as well as the hidden state of the last time. Due to the long-term dependence on the gradient of the hidden layers, one of the obvious flaws of RNN is gradient disappearance and explosion. Hence, gated recurrent unit (GRU) and long short-term memory (LSTM) are developed (Cho *et al.*, 2014; Hochreiter and Schmidhuber, 1997). Compared with traditional RNN neurons, gated systems are added to these two RNN variants, which can better overcome the drawbacks of gradients disappearance and explosion mentioned above. Moreover, to improve the prediction accuracy, we propose bidirectional GRU (BGRU) and bidirectional LSTM (BLSTM), which can use the past and future information from the current time as input (Zhang *et al.*, 2015). Furthermore, we integrated the attention mechanism with BGRU and BLSTM modules to improve the model (Zhou *et al.*, 2016). As the name implies, the attention mechanism focuses limited attention on key information and adjusts the model according to specific task objectives to make sequence-based modelling easier.

**Model training and validation**

Predictive models were implemented in Python based on the Pytorch framework. Binding affinity is converted to a value between 0 and 1 according to equation (1). The encoding matrix along with the corresponding labels are as the input, and one of the six RNN variants (GRU, LSTM, BGRU, BLSTM, att-BGRU, att-BLSTM) was chosen for training respectively. After experimental verification (Table. S1), the three layers of BGRU are adopted as hidden layers, sigmoid is used as the activation function of the output layer, the binary cross-entropy and mean-square loss (MSE) are respectively employed to calculate the loss in classification and regression task, and parameters are optimized using an Adam optimizer with a default learning rate of 0.001. About 10% of the data is preserved for subsequent model testing, and five-fold cross-validation is used to train the network on the remaining data. The dataset is randomly divided into five subsets according to the proportion of different label values. The model is trained on four subsets while tested on the fifth subset and repeated five times. The final prediction results represent the average value predicted by five independent neural networks. For the binary classification model, the predicted score is between 0 and 1, more than 0.5 is considered a TAP binding peptide. For the regression model, the prediction score is the log value of relative IC50.

$$value = 1 - log_{50000000}(nM\ affinity) \qquad (1)$$

**Comparison with traditional SVM and RF**

We expanded the dataset for model training based on the previous, while compared with other machine learning tasks, the sample dataset is still relatively small. We wondered whether the traditional machine learning algorithms can achieve more accurate prediction, therefore support vector machine (SVM) based radial basis function (RBF) and random forest (RF) are employed for model training and testing, and compared with RNN. For the classifier, the area under the receiver operating characteristic curve (AUC) is the main measurement. For the classification task, DeepTAP, BGRU-based, achieves the best prediction performance with an AUC of 0.95, following is RF with 0.9 and SVM with 0.8 (Table. S2). For the regression task, the parallel performance was obtained in Table. S3. Results show that the neural network based on RNN can obviously improve the prediction accuracy compared with the traditional machine learning algorithm, and may be a more reliable method for predicting the binding between peptide and TAP.

**Comparison with other available tools**

Moreover, to verify the reliability of our model, other available prediction tools including TAPPred (Bhasin and Raghava, 2004), TAPREG (Diez-Rivero *et al.*, 2010) and KSMM (Hao *et al.*, 2020) are employed to compare with the RNN-based DeepTAP. Other algorithms are excluded from evaluation due to challenging to use interfaces (Peters *et al.*, 2003; Zhang *et al.*, 2006; Ren *et al.*, 2011). TAPPred is an online TAP binding peptide prediction tool developed by Bhasin and Raghava et al based on simple SVM and cascade SVM. Two-layer SVM is used in cascade SVM, where the first layer includes 33 SVMs, which combines amino acid sequence with the information of physical and chemical properties as input, involving 33 features, and then takes the output of the first layer as the input of the second layer to get a final prediction score. TAPREG is a prediction tool based on SVM, and KSMM is a scoring matrix algorithm with kernel function, which takes into account the pairwise interaction of amino acids at different positions of the peptide sequence. All of them obtain great prediction performance in their own published papers. The benchmark evaluation is only carried out on the classification task. For the regression task, it cannot be directly compared with the existing tools, the evaluation results of each tool are directly quoted. Though data in the benchmark dataset may overlap with the training data of previous software, DeepTAP still performs much better than all other software with the highest AUC of 0.95 (Table. S2). Similar prediction results have been achieved in several methods based on SVM with the approximate AUC of 0.8, while the prediction performance of cascade SVM and KSMM are the worst. One of the most likely reasons is that after combining the physical and chemical properties or interaction of amino acids with sequences, the input information is so complex that the model cannot extract and learn correct feature information, resulting in poor training and prediction results. Another measurement, accuracy (ACC) is likewise usually employed to evaluate the quality of the classification system. Similarly, the results show that the ACC value of the RNN-based model is higher than others, in addition to AUC and ACC, AP, sensitivity and specificity are also often used to evaluate the prediction performance of classification methods. Parallel performance is obtained in the regression task (Table. S3), where the model performance is evaluated by the Spearman correlation coefficient (Spearman r) and Pearson correlation coefficient (Pearson r). Therefore, a more detailed analysis has been carried out in this paper, and the results show that RNN-based DeepTAP is superior to the existing prediction tools in terms of prediction accuracy and error control.

### Supplementary Figures

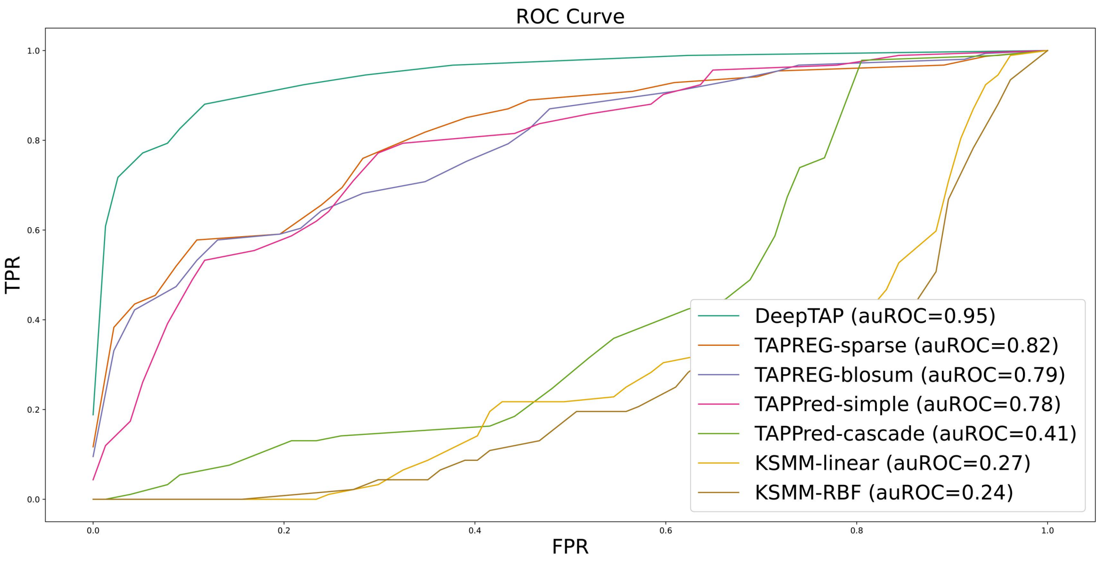


**Figure S1.** Comparison of ROC curves for RNN, SVM, RF and other methods on the benchmark dataset, showing the AUC value of DeepTAP (our tool, 0.95), TAPREG-sparse (0.82), TAPREG-blosum (0.79), TAPPred-simple (0.78), TAPPred-cascade (0.41), KSMM-linear (0.27) and KSMM-RBF (0.24).

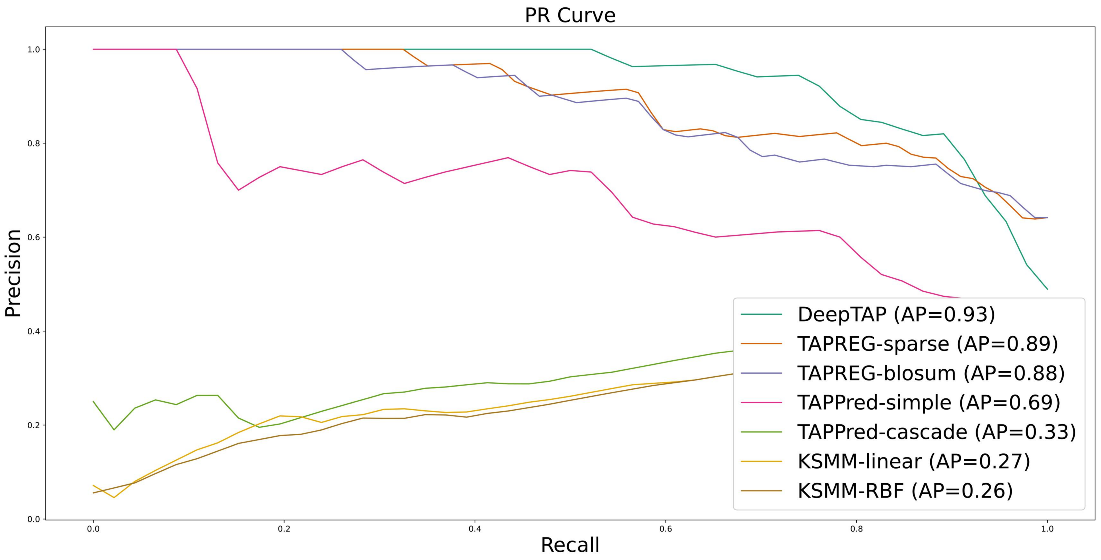


**Figure S2.** Comparison of PR curves for RNN, SVM, RF and other methods on the benchmark dataset, showing the AP value of DeepTAP (our tool, 0.93), TAPREG-sparse (0.89), TAPREG-blosum (0.88), TAPPred-simple (0.69), TAPPred-cascade (0.33), KSMM-linear (0.27) and KSMM-RBF (0.26).

### Supplementary Tables

**Table S1. RNN-based model performance with cross-validation**

| Model | classifier | | | | | regression | |
|---|---|---|---|---|---|---|---|
| | AUC | AP | ACC | Sensitivity | Specificity | Spearman r | Pearson r |
| GRU | 0.89 | 0.81 | 0.82 | 0.83 | 0.80 | 0.81 | 0.82 |
| LSTM | 0.88 | 0.80 | 0.80 | 0.78 | 0.83 | 0.75 | 0.72 |
| **BGRU** | **0.94** | **0.93** | **0.85** | **0.82** | **0.88** | **0.89** | **0.88** |
| BLSTM | 0.90 | 0.84 | 0.84 | 0.80 | 0.87 | 0.87 | 0.87 |
| att_BGRU | 0.90 | 0.85 | 0.83 | 0.78 | 0.85 | 0.86 | 0.85 |
| att_BLSTM | 0.90 | 0.84 | 0.82 | 0.82 | 0.84 | 0.82 | 0.81 |

**Table S2. Classification model comparison on benchmark dataset**

| Model | AUC | AP | ACC | Sensitivity | Specificity |
|---|---|---|---|---|---|
| **DeepTAP** | **0.95** | **0.93** | **0.87** | **0.80** | **0.91** |
| RF | 0.90 | 0.85 | 0.92 | 0.83 | 0.97 |
| SVM | 0.80 | 0.71 | 0.84 | 0.65 | 0.95 |
| TAPREG_sparse | 0.82 | 0.89 | 0.76 | 0.72 | 0.79 |
| TAPREG_blosum | 0.79 | 0.88 | 0.72 | 0.61 | 0.78 |
| TAPPred_simple | 0.78 | 0.69 | 0.72 | 0.80 | 0.66 |
| TAPPred_cascade | 0.41 | 0.33 | 0.47 | 0.98 | 0.17 |
| KSMM_linear | 0.27 | 0.27 | 0.30 | 0.37 | 0.26 |
| KSMM_rbf | 0.24 | 0.26 | 0.30 | 0.35 | 0.27 |

**Table S3. Regression model comparison on benchmark dataset**

| Model | Spearman r | Pearson r | Reference |
|---|---|---|---|
| **DeepTAP** | **0.90** | **0.88** | **This study** |
| SVM | 0.72 | 0.72 | This study |
| RF | 0.82 | 0.82 | This study |
| TAPREG | 0.89 | - | (Diez-Rivero et al., 2010) |
| TAPPred | 0.88 | - | (Bhasin and Raghava, 2004) |
| SVMTAP | 0.82 | - | (Dönnes and Kohlbacher, 2005) |
| ADM | 0.74 | - | (Doytchinova *et al.*, 2004) |
| SMM | 0.82 | - | (Peters *et al.*, 2003) |
| KSMM | 0.89 | - | (Hao *et al.*, 2020) |
